# Comparison of oral and gut microbiome highlights role of oral bacteria in systemic inflammation in HIV

**DOI:** 10.1101/2025.05.16.654362

**Authors:** Jennifer A. Fulcher, Katharine Newman, Bryan Pham, Fan Li, Grace D. Cho, Julie Elliott, Nicole H. Tobin, Steven Shoptaw, Pamina M. Gorbach, Grace M. Aldrovandi

**Affiliations:** Division of Infectious Diseases, Department of Medicine, David Geffen School of Medicine at UCLA; Division of Infectious Diseases, Department of Pediatrics, David Geffen School of Medicine at UCLA; Department of Family Medicine, David Geffen School of Medicine at UCLA; Department of Epidemiology, Fielding School of Public Health, University of California, Los Angeles

## Abstract

**Background:** Chronic HIV-1 infection is associated with increased inflammation-related comorbidities, despite effective viral suppression with antiretroviral therapy. While the role of the gut microbiome in inflammation is well-studied, the contribution of the oral microbiome remains less clear. This study investigates the relationship between the oral and gut microbiomes in driving systemic inflammation in persons with HIV.

**Methods:** This cross-sectional study utilized archived samples from 198 participants (99 with HIV and 99 without HIV). Oral microbiome composition was analyzed via 16S rRNA sequencing and systemic inflammatory biomarkers were measured using multiplex assays. Gut microbiome data from previous studies were integrated for comparative analyses. Bacterial inflammatory potential was assessed through in vitro co-culture and epithelial barrier permeability assays.

**Results:** The oral microbiome in HIV was characterized by increased *Veillonella, Capnocytophaga*, and *Megasphaera*, and several decreased genera including *Fusobacterium*. Using PERMANOVA, we found that the oral microbiome was a significant driver of cytokine variation in HIV compared to the gut microbiome, and identified specific associations with oral *Veillonella* and *Megasphaera*. We found no differences in anti-*Veillonella parvula* serum IgG by HIV status, but IgG titers did correlate with microbial translocation markers sCD14 and LBP in HIV. *In vitro* studies demonstrated that *Veillonella parvula* increased oral epithelial barrier permeability and induced monocyte activation.

**Conclusions:** The oral microbiome, particularly *Veillonella parvula*, may contributes to systemic inflammation in HIV through mechanisms involving epithelial barrier disruption, oral translocation, and monocyte activation.

## INTRODUCTION

Chronic HIV-1 infection is characterized by accelerated risk of inflammation-related comorbidities. Despite the ability of antiretroviral therapy to effectively control viremia, persons living with HIV (PWH) have increased risk of non-AIDS related comorbidities including cardiovascular disease and neurocognitive disease. While the cause is likely multifactorial, ample data supports the role of inflammation in these processes^[1-4]^, which is driven by mucosal barrier dysfunction and microbial translocation^[5-7]^.

While the contribution of the gut microbiome to chronic inflammation in HIV is well-described^[5, 8]^, less is known about the role of the oral microbiome in inflammation in PWH. While several studies have found associations between HIV and oral dysbiosis^[9-16]^, there is significant variation across studies due to variations in study design, sample size, and sampling techniques. Findings reported across multiple studies include alterations in *Veillonella, Fusobacterium, Rothia, Neisseria* and *Streptococcus*^[10, 11, 13-17]^. Several studies have found conflicting results regarding changes over time during HIV^[14, 18]^, as well as the impact of antiretroviral therapy on the oral microbiome^[10, 13, 17, 19]^.

Some studies have examined associations between the oral microbiome and inflammation in HIV. Yang et al found that *Veillonella, Streptococcus*, and *Lactobacillus* all associated with inflammatory biomarkers, and were further associated with decreased lung function^[15]^. At least one study simultaneously examined the oral and gut microbiomes in PLWH and found that systemic inflammation biomarkers were inversely associated with saliva bacterial diversity, but not associated with the gut microbiome, suggesting a role for the oral microbiome in systemic inflammation in HIV^[12]^.

The oral microbiome has been associated with multiple systemic diseases including cardiovascular disease, inflammatory bowel disease, rheumatoid arthritis, and dementia^[20]^. Translocation of oral bacteria, primarily via the bloodstream, is one mechanism underlying these associations. Oral commensal bacteria, including *Fusobacterium nucleaum* and *Streptococcus spp*., have been identified in atherosclerotic plaques in coronary artery disease^[21]^, suggesting a primary role for these bacteria in the bloodstream in cardiovascular disease. Ectopic colonization of the gastrointestinal tract by oral bacteria has also been implicated in disease development. Ectopic colonization of *Veillonella parvula* in the gut promoted development of colitis in a mouse model^[22]^. We hypothesize that similar mechanisms contribute to inflammation in HIV. Here we examine the relative contribution of the oral and gut microbiome to systemic inflammation in HIV, and test the inflammatory potential of key bacteria.

## METHODS

### Study design and specimens

This cross-sectional study used archived specimens from the mSTUDY cohort (U01DA036267), a longitudinal cohort of men who have sex with men with and without HIV including half who actively use drugs based on Los Angeles, CA. The mSTUDY was approved by the UCLA Office of the Human Research Protection Program Institutional Review Board (IRB) and all participants signed written informed consent at the time of enrollment. Participants in the mSTUDY attend study visits approximately every six months which consist of collection of clinical data including laboratories, stored peripheral blood mononuclear cells, plasma, and serum, urine drug screen, stored rectal swab, stored saliva and oral rinse, and complete computer assisted self-interview (CASI) questionnaires detailing sexual behavior and substance use. The current study included blood and saliva samples from 198 mSTUDY participants selected based on paired blood and saliva sample availability from the first study visit. mSTUDY samples used in this study were collected between 2014 and 2018. Gut microbiome data obtained from the same participants and study visits in prior studies^[23, 24]^ were also included in select analyses. Demographic and behavioral data was obtained from cohort study records. Saliva samples were collected by passive drool method then frozen at - 80°C until processing in bulk.

### Oral microbiome analysis

Saliva processing and sequencing was performed by the University of California at San Diego (UCSD) Microbiome Core as previously described^[25]^. Briefly, DNA extraction was performed using the MagMAX Microbiome Ultra Nucleic Acid Isolation Kit (Thermo Fisher Scientific) followed by sequencing of the V4 region of the 16S rRNA on the Illumina NovaSeq 6000 platform (2×150). Negative controls (extraction blanks and PCR negatives) and positive controls (ZymoBIOMICS Microbial Community Standard) were sequenced concurrently. Dada2 (v1.22.0), decontam (v.1.14.0), and phyloseq (v1.38.0) were used for sequence inference, contaminant removal, and subsequent analyses. Taxonomic assignment was performed using the RDP training set version 18 and the “assignTaxonomy” and “addSpecies” functions from the “dada2” package with default parameters.

### Serum cytokine quantification

Serum samples from the same participants and study visits were used for cytokine analyses. Serum cytokines/biomarkers were measured using Meso Scale multiplex assays (Meso Scale Diagnostics) and ELISAs (R&D Systems) per manufacturer’s instructions. All samples were run in duplicate and processed in bulk to reduce inter-assay variability. Samples with a mean coefficient of variation (%CV) >20% were repeated. Values below the lower limit of quantification were imputed as LLOQ/2.

### Bacteria culture and LPS purification

*Fusobacterium nucleatum subsp. nucleatum* (ATCC 25586) and *Veillonella parvula* strain [ATCC 17742, Te 3] were purchased from American Type Culture Collection (Manassas, VA, USA). Bacteria were grown under anaerobic conditions at 37°C in thioglycollate broth (cat#AS-801, Anaerobe Systems). Cultures were grown to OD_600_ 0.8-1.2 then lipopolysaccharide (LPS) was purified using iNtRON LPS Extraction Kit according to manufacturer’s instructions (Boca Scientific). Purified LPS was quantified using the Pierce Chromogenic Endotoxin Quant Kit according to manufacturer’s instructions (cat# A39553, Thermo Scientific).

### Anti-bacteria IgG ELISA

Bacteria were grown as described above for 24h-48h (OD_600_ 0.8-1.2) then cells harvested by centrifugation at 10,000xg for 10 minutes. Cell pellet was lysed using BugBuster Protein Extraction Reagent (Novagen) then debris removed by centrifugation at 20,000xg for 10 minutes. Supernatant was collected then protein concentration measured using Pierce BCA Protein Assay. 96-well MaxiSorp plates (Nunc) were coated overnight with 50ul of 0.5mg/ml bacteria lysate. Plates were then washed with PBS/0.1%Tween (TPBS) and blocked with 3% milk/TPBS for 2 hours at room temperature. Plates were washed with TPBS then serum samples were added in serial dilutions from 1:500 to 1:13,500 for 2 hours at room temperature. After washing with TPBS, goat anti-human IgG-HRP (Bethyl Laboratories) 1:25,000 was added for 1 hour at room temperature. Plates were again washed then TMB substrate added followed by stop solution at 20 minutes. Absorbance was measured at OD450 and OD650 using plate reader (Tecan Spark). Endpoint titers were interpolated as described^[26]^. Pre-tested negative human serum was used to determine positive cutoff set at two standard deviations above the mean OD of negative serum. Interpolated endpoints were then compared by HIV serostatus using standard non-parametric tests (Mann-Whitney). All analyses were performed using Prism v10.4.2 (GraphPad).

### PBMC co-cultures

Bacteria were grown as described above for 24h-48h (OD_600_ 0.8-1.2) then cells harvested by centrifugation at 10,000xg for 10 minutes. Bacteria were resuspended in PBS then heat-inactivated by incubation at 56°C for 30 minutes. Bacteria were counted by flow cytometry using Bacteria Counting Kit with SYTO BC dye (cat#B7277 Invitrogen). PBMC were co-cultured in RPMI with 10% FBS overnight with 10ng/ml LPS or inactivated bacteria at a bacteria to cell ratio of 25:1. Following co-culture PBMC were washed and stained for flow cytometry with the following antibodies: Pacific Blue anti-human CD3 (BioLegend cat#300431, clone UCHT1), Brilliant Violet anti-human CD16 (BioLegend cat#302040, clone 3G8), Brillian Violet anti-human CD11c (BioLegend cat#337238, clone Bu15), BB515 anti-human HLA-DR (BD Horizon cat#564516, clone G46-6), PE anti-human CD38 (BD Pharmingen cat#555460, clone HIT2), PE/Dazzle 594 anti-human CD19 (BioLegend cat#302252, clone HIB19), PE-Cy7 anti-human CD4 (BioLegend cat#344612, clone SK3), APC anti-human CD14 (BioLegend cat#301808, clone M5E2), APC-H7 anti-human CD8 (BD Pharmingen cat#560179, clone SK1). Data was acquired on a BD FACSymphony A1 cell analyzer. Flow cytometry data were analyzed using FlowJo (v10.10.0).

### Oral epithelial barrier assay

TR146 buccal carcinoma epithelial cell line was purchased from Sigma-Aldrich (cat#10032305). and grown to confluency in a transwell (Corning 24-well 0.4μm pore) in DMEM with 10% FBS. Cells were plated at 24,000 cells/cm2 in a transwell and cultured with media exchange every 2-3 days until confluent (total culture 1-2 weeks). The integrity and barrier function of the confluent transwells was confirmed by measuring the transepithelial electrical resistance (TEER) using a dual electrode connected to an epithelial volt/ohm meter (World Precision Instruments, Sarasota, FL). Confluent transwells were then treated overnight with bacteria (25:1 bacterial to cell ratio) or LPS (10ng/ml) or media in the apical chamber then barrier function assessed using dextran flux. After treatment with bacteria/LPS, the apical medium was removed and replaced with DMEM without FBS for 30 minutes at 37°C. FITC-Dextran (4kDa, Sigma cat#46944) was added to the apical chamber (1mg/ml) then 100μl media from the bottom chamber was sampled immediately and every 30 minutes for 2 hours. Fluorescence was measured using Tecan Spark fluorimeter (485nm excitation, 525 emission). A control transwell with no cells was included as a measure of maximum flux, and all conditions were normalized as a percent of maximal flux for that time point (percent permeability). The percent barrier permeability for each condition was compared using Friedman tests. All analyses were performed using Prism v10.4.2 (GraphPad).

### Statistical analyses

Statistical analyses of the microbiome data were performed using the phyloseq (v1.38.0), vegan (v2.6.2), glmmTMB (v1.1.6), emmeans (v1.8.1.1), and twang (v2.5) packages within the R statistical environment (v4.1.2). Propensity scores were calculated using gradient boosted logistic regression and used in as weights in the subsequent regression models. An offset term was also applied based on geometric mean of pairwise ratios (GMPR) normalization. Alpha diversity metrics were calculated using a rarefaction of 50318 reads, and differences were assessed using linear regression models with estimated marginal means. To test for specific taxonomic differences between groups, we first fit negative binomial regression models both with and without zero inflation for each taxon. The better model for each taxon was then selected according to the Akaike information criterion (AIC). To quantify the cytokine variation attributable to oral and gut microbiome profiles, we used a PERMANOVA-based approach. Specifically, we first computed the first 20 principal components for each microbiome compartment. PERMANOVA was then used to identify PCs that contributed significantly (p<0.1) to the plasma cytokine profile. Finally, the percentage of cytokine variation attributable to each compartment was calculated as the sum of the variation attributable to all significant PCs from the given compartment. Integrative analysis of oral and gut microbiome profiles and plasma cytokines was performed using a multiblock sparse partial least squares discriminant analysis approach as implemented by the DIABLO method in the mixOmics R package (v6.25.1). For all other experiments, standard statistical tests were used as noted in the Methods and Figure Legends.

## RESULTS

### Study design and sample characteristics

This study included samples from 198 individuals including 99 persons with HIV (PWH) and 99 persons without HIV (PWOH). Participant demographics and characteristics are shown in **Table 1**. All participants were men who have sex with men. The group with HIV was older (34 vs 30 years; p<0.001). Among those with HIV, the majority had well-controlled HIV (80.8% with viral load <200 copies/ml) and all participants had CD4 T cells >200 cells/ml. There were no significant differences by HIV status in terms of racial/ethnic groups, employment status, or body mass index. Sexual behavior was similar between those with and without HIV. The majority of participants did not smoke and had infrequent binge alcohol use, with no significant differences between those with and without HIV. Almost half of participants used methamphetamine, with no significant differences by HIV status in terms of frequency. More participants without HIV used cannabis than with HIV (53.5% vs 25.3%; p<0.0001).

**Table 1.** Study participant demographics and characteristics

|  | Without HIV<br>(n=99) | With HIV<br>(n=99) | *p-value*<sup>a</sup> |
| --- | --- | --- | --- |
| **Age** (mean (sd), median) | 30 (7.2), 28 | 34 (6.3), 35 | <0.001 |
| **Race/Ethnicity** (n,%) |  |  | 0.15 |
| Black-Non Hispanic | 43 (43.4) | 30 (30.3) |  |
| Hispanic | 45 (45.5) | 57 (57.6) |  |
| Other-Non Hispanic | 11 (11.1) | 12 (12.1) |  |
| **Employment** (n,%) |  |  | 0.30 |
| Full/part time | 48 (48.5) | 42 (42.4) |  |
| Unemployed | 35 (35.4) | 42 (42.4) |  |
| Disabled | 6 (6.0) | 10 (10.1) |  |
| Student | 10 (10.1) | 5 (5.1) |  |
| **Body mass index (BMI)** (mean (sd), median) | 28 (6.5), 26 | 27 (5.6), 26 | 0.62 |
| **Viral load** (n,%) |  |  | NA |
| (log mean (sd), median) |  |  |  |
| Undetectable (<20 copies/ml) | -- | 42 (42.4) |  |
|  |  | -- |  |
| Suppressed (<200 copies/ml) | -- | 38 (38.4) |  |
|  |  | 3.4 (0.65), 2.9 |  |
| Unsuppressed (>200 copies/ml) | -- | 19 (19.2) |  |
|  |  | 9.1 (2.4), 9.8 |  |
| **CD4 T cells** (mean (sd), median) | -- | 667 (288), 638 | NA |
| **Antiretroviral therapy** (n, %) |  |  | NA |
| NNRTI | -- | 33 (35.9) |  |
| PI | -- | 17 (18.5) |  |
| INSTI | -- | 42 (45.6) |  |
| **Number of oral sex acts** (30 days)<br>(mean (sd), median) | 12 (24), 5 | 14 (30), 5 | 0.89 |
| **Number of RAI acts** (30 days) (mean<br>(sd), median) | 2.3 (5.1), 1 | 3.6 (6.9), 1 | 0.22 |
| **Number of sex partners** (30 days)<br>(mean (sd), median) | 6.9 (8.5), 4 | 8.1 (9.3), 5 | 0.61 |
| **Methamphetamine use**<sup>c</sup> (n,%) |  |  | 0.16 |
| Daily | 11 (11.1) | 18 (18.2) |  |
| Weekly | 7 (7.1) | 13 (13.1) |  |
| Monthly/less | 27 (27.3) | 19 (19.2) |  |
| Never | 54 (54.5) | 49 (49.5) |  |
| **Cannabis use** (n,%) |  |  | <0.0001 |
| Daily | 11 (11.1) | 0 (0) |  |
| Weekly | 8 (8.1) | 6 (6.1) |  |
| Monthly/less | 34 (34.3) | 19 (19.2) |  |
| Never | 46 (46.5) | 74 (74.7) |  |
| **Smoking** (n,%) |  |  | 0.61 |
| >1 pack/day | 7 (7.0) | 4 (4.0) |  |
| <1 pack/day | 37 (37.4) | 36 (36.4) |  |
| Nonsmoker | 55 (55.6) | 59 (59.6) |  |
| **Binge alcohol use**<sup>c</sup> (n,%) |  |  | 0.14 |
| Weekly | 12 (12.1) | 12 (12.1) |  |
| Monthly/less | 46 (46.5) | 33 (33.3) |  |
| Never | 41 (41.4) | 54 (54.6) |  |

<sup>a</sup>t-tests or Mann-Whitney tests were used for continuous variables and Chi-squared tests for categorical variables
<sup>b</sup>sd, standard deviation; RAI, receptive anal intercourse; NA, not applicable
<sup>c</sup>Substance use data refers to past 6 month and was collected by self-report. Binge alcohol refers to >6 drinks at a time on more than 1 occasion.

### Oral microbiome composition differs between persons with and without HIV

The relative oral microbiome composition for each participant is shown in **Figure 1A**. The most common genera in both groups include *Haemophilus, Veillonella, Streptococcus*, and *Prevotella*. There were no significant differences in alpha diversity between HIV groups (**Figure 1B**), however, principal coordinates analysis did show differences between those with HIV and without HIV (**Figure 1C**). We used generalized linear models to identify specific genera that distinguish the oral microbiome between PWH and PWOH. All analyses were adjusted for multiple confounders (age, substance use, sexual behavior, viral load, etc) using a propensity-score based method as previously described^[23, 27]^. The oral microbiome in PWH showed decreased *Lactobacillus, Lautropia, Treponema, Oribacterium*, and *Fusobacterium*, as well as increased *Veillonella, Capnocytophaga*, and *Megasphaera* (**Figure 1D and Supplementary Table 1**) compared to PWOH.

**Figure 1.**
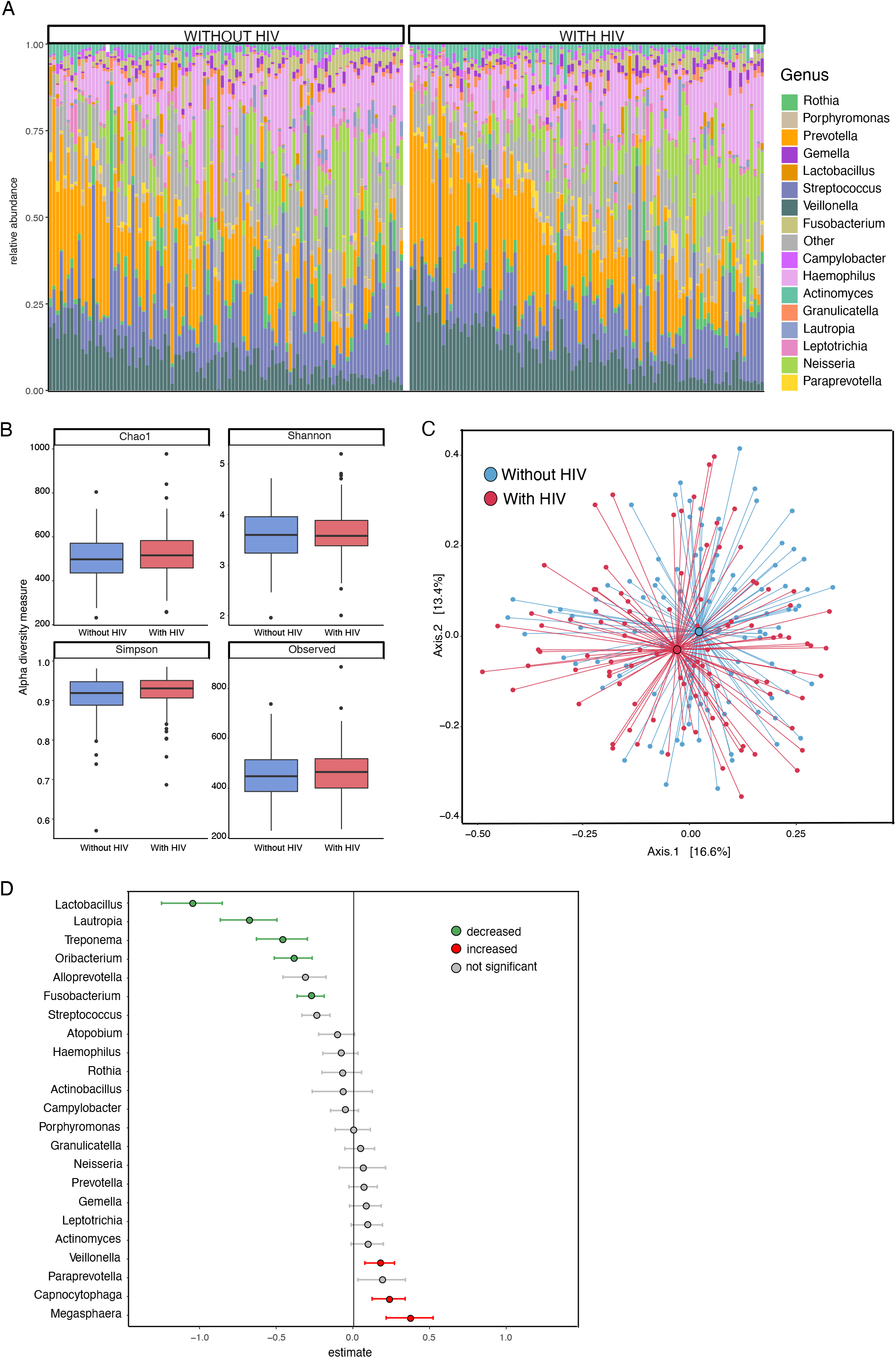
Oral microbiome composition differs in HIV. **(A)** Taxa barplot showing oral microbiome composition for participants without HIV (left) and with HIV (right). Each bar represents individual participants and colors show averaged relative abundance as a percentage of total bacterial sequences at the genus level. **(B)** Oral microbiome alpha diversity measures are not significantly different between persons with (red) and without (blue) HIV. **(C)** Principal coordinates analysis showing differences in oral microbiome between persons with (red) and without (blue) HIV. **(D)** Forest plot showing specific bacterial genera increased or decreased with HIV using zero-inflated negative binomial regression analysis. Green or red indicates associations that remained statistically significant (q < 0.05) after FDR-adjustment.

### Oral microbiome contributes significantly to systemic cytokine variation

Plasma cytokines and immune biomarkers were compared between PWH and PWOH. There were significant increases in several inflammatory cytokines/biomarkers in PWH, including TNF-α, IL-6, sCD163, sCD27, CXCL10, and FABP2 (**Supplementary Figure 1**). Using gut microbiome data from prior studies on the same participants and study visits^[23, 24]^, we quantified the contribution of each microbiome compartment on plasma cytokine variation using PERMANOVA. As shown in **Figure 2A**, the oral microbiome is a significant contributor to cytokine variation. We then used integrative analytic approaches to identify relationships between the two microbiome compartments and plasma cytokines. **Figure 2B** shows that both oral *Veillonella* and *Megasphaera* were positively correlated with IL-6 and FABP2, suggesting a potential role in systemic inflammation. Additionally, both oral *Veillonella* and *Megasphaera* were negative correlated with gut *Bacteroides*, which may suggest a role in gut dysbiosis. Finally, gut *Porphyromonas* was positive correlated with sCD27.

**Figure 2.**
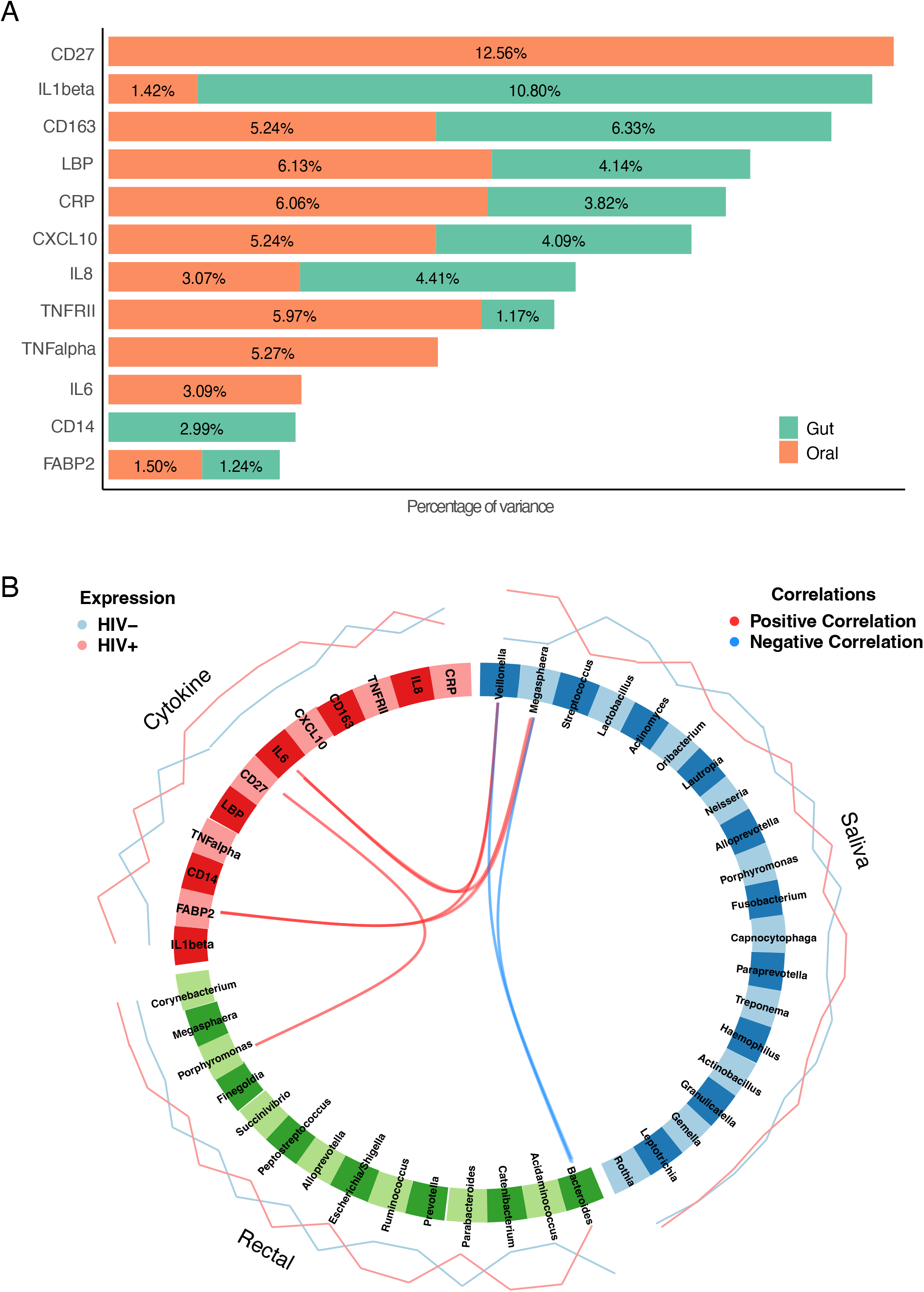
Relative contributions of oral and gut bacteria to cytokine variation in HIV. **(A)** Relative contribution of the gut (green) or oral (orange) microbiome to serum cytokine variation as determined by PERMANOVA. (B) Circos plot showing positive correlations (red) between *Porphyromonas* in the gut and *Veillonella* and *Megasphaera* in the saliva with inflammatory cytokines, and negative correlations (blue) with gut *Bacteroides*.

### Plasma anti-bacteria IgG does not differ by HIV status

Given the positive correlation between *Veillonella* and *Megasphaera* with inflammatory cytokines, we hypothesized that systemic circulation of these bacteria could be a trigger for inflammation. To look for evidence of systemic circulation, we quantified anti-bacteria IgG in the plasma. For these two genera, we examined the species-level data (**Supplementary Table 1**) and identified *Megasphaera micronuciformis* and *Veillonella parvula* as the most significant species in each genus. We focused on *Veillonella parvula*, and used *Fusobacterium nucleatum* as representative of decreased genera based on availability. There were no differences in anti-*Veillonella parvula* IgG or anti-*Fusobacterium nucleatum* IgG in PWH compared to PWOH (**Figure 3A**).

**Figure 3.**
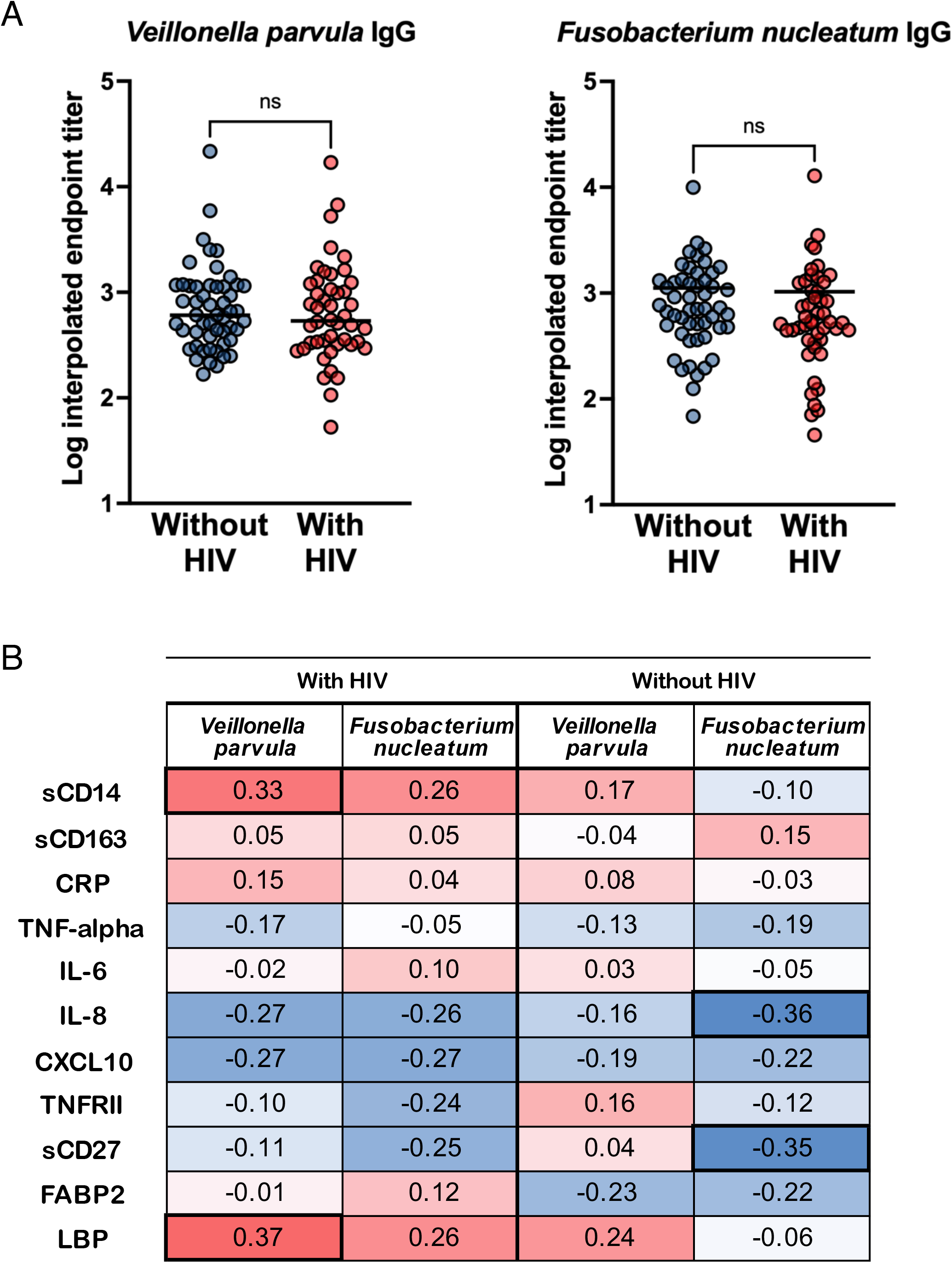
Anti-bacteria IgG endpoint titers correlate with microbial translocation markers in HIV. **(A)** Anti-*Veillonella parvula* or anti-*Fusobacterium nucleatum* IgG endpoint titers were measured in plasma by ELISA. Endpoint titers were interpolated from serial dilutions using negative pooled plasma as cutoff. Titers were compared between HIV serogroups using Mann-Whitney tests. **(B)** Spearman correlation between inflammatory cytokines/biomarkers and anti-bacteria endpoint titers. Heatmap shows results separated by HIV status. Numbers denote Spearman rho values with significant values (p<0.05) in bold.

### Anti-*Veillonella parvula* IgG correlates with microbial translocation markers in HIV

We next examined the relationship between anti-bacteria IgG and plasma cytokines using multiple linear regression (**Supplementary Table 2**). As expected, HIV was a significant predictor for several cytokines (sCD14, LBP, sCD163, FABP2, IL-6, TNF-alpha, sCD27, CXCL10). *Veillonella parvula* IgG was a significant predictor for sCD14 (β=0.17, p=0.01) and LBP (β=0.18, p=0.02), and *Fusobacterium nucleatum* IgG was a significant predictor for IL-8 (β=-0.14, p=0.03) and sCD27 (β=-0.11, p=0.01). Similarly, we examined Spearman correlations and again found significant positive correlation between *Veillonella parvula* IgG and microbial translocation markers sCD14 (r=0.33, p=0.03) and LBP (r=0.37, p=0.02), but only in PWH (**Figure 3B**). *Fusobacterium nucleatum* IgG negatively correlated with IL-8 (r=-0.36, p=0.01) and sCD27 (r=-0.35, p=0.01), but only in PWOH.

### *Veillonella parvula* increases oral epithelial barrier permeability

To further examine the hypothesis that *Veillonella parvula* oral translocation could contribute to systemic inflammation in HIV, we examined the effects of *Veillonella parvula*, both whole bacteria and isolated LPS, on oral epithelial barrier function. Using TR146 cells in a transwell system^[28]^, we found that *Veillonella parvula* LPS, but not LPS from other bacteria, increased epithelial barrier permeability (**Figure 4A**).

**Figure 4.**
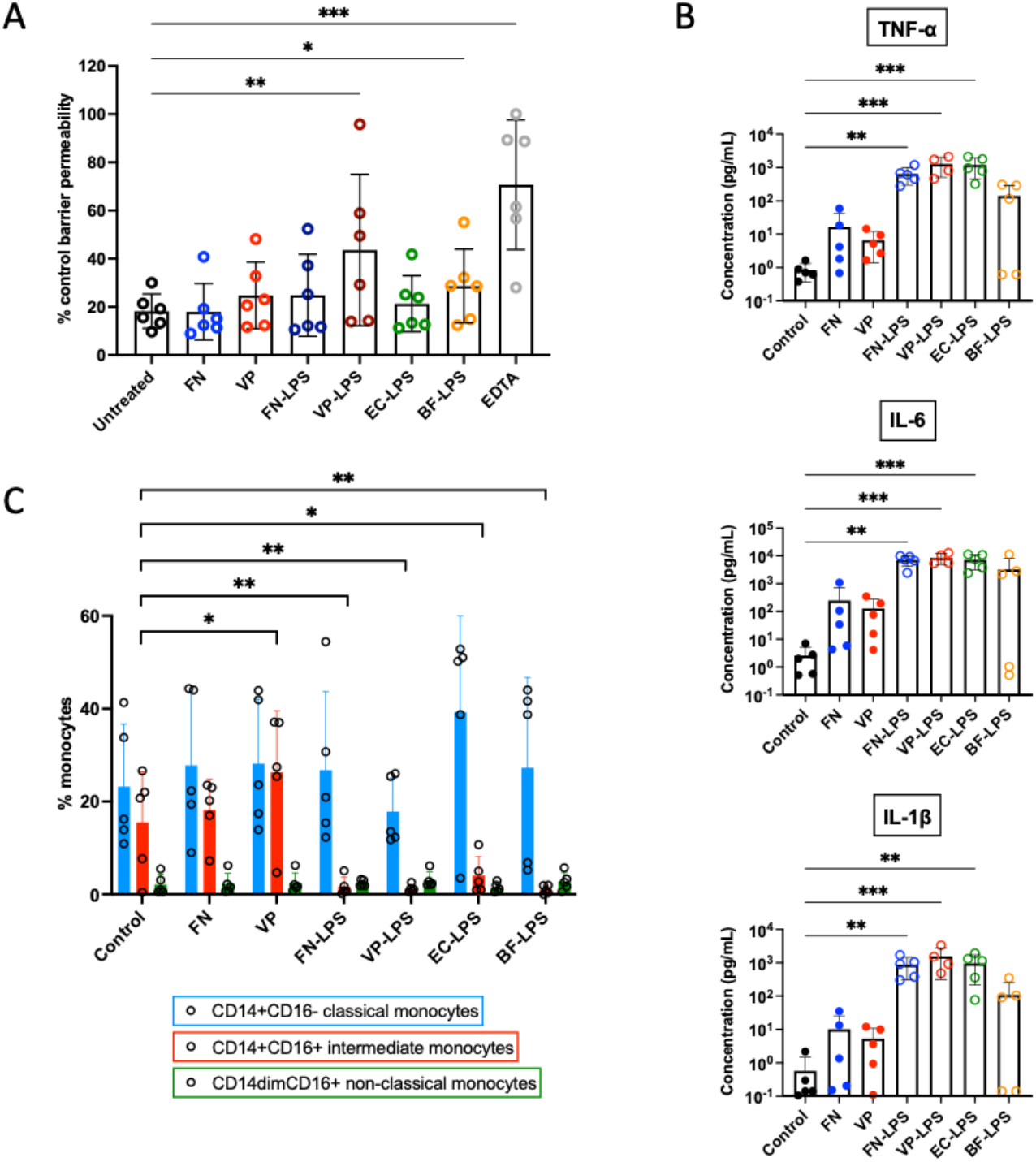
Veillonella parvula alters oral epithelial barrier function and induces PBMC responses. **(A)** FITC-dextran permeability assay using TR146 cells shows increased permeability following treatment with either heat-inactivated bacteria or isolated LPS from *Veillonella parvula* (VP). Permeability expressed as percent of transwell flux with no cells (set at 100%). Bars shown mean with error bars showing standard deviation from six independent experiments. * p<0.05; ** p<0.01; ***p<0.001 by Friedman tests. FN, *Fusobacterium nucleatum*; EC, *Escherichia coli*; BF, *Bacteroides fragilis*. **(B)** Increased inflammatory cytokines production in supernatant from PBMC co-cultures with heat-inactivated bacteria or isolated LPS. Bars shown mean with error bars showing standard deviation from five independent donors. ** p<0.01; ***p<0.001 by Kruskal-Wallis tests. **(C)** Monocyte subsets as a percentage of total monocytes following co-culture with heat-inactivated bacteria or isolated LPS. Bars shown mean with error bars showing standard deviation from five independent donors. * p<0.05; ** p<0.01 by one-way ANOVA.

### *Veillonella parvula* induces pro-inflammatory cytokine production and monocyte activation

To examine the capacity for *Veillonella parvula* to induce peripheral inflammation we performed co-culture experiment with peripheral blood mononuclear cells (PBMC). We found that LPS from *Veillonella parvula* significantly increased pro-inflammatory cytokine production, similar to LPS from *Fusobacterium nucleatum* and *Escherichia coli* (**Figure 4B**). Co-culture with *Veillonella parvula* bacteria, but not LPS, increased the proportion of the inflammatory intermediate (CD14+CD16+) monocytes subset (**Figure 4C**). There were no changes in other cell populations activation, such as T cells (data not shown).

## DISCUSSION

In this study we identified several alterations to the oral microbiome in HIV, including increased *Veillonella, Capnocytophaga*, and *Megasphaera*. Using paired gut microbiome data, we also showed the significant contribution of the oral microbiome, particularly *Veillonella parvula*, to systemic inflammation in PWH.

Consistent with previous studies, our analysis revealed distinct differences in the oral microbiome composition between PWH and PWOH. We observed increased *Veillonella, Capnocytophaga*, and *Megasphaera* in HIV, along with a reduction in *Fusobacterium* among others (**Figure 1**). Previous studies reported similar shifts in oral microbiota, specifically increased *Veillonella* and/or *Megasphaera*^[9, 11, 13]^. Alterations in *Fusobacterium* have also been reported in HIV^[12]^, though many other studies reported increases while we showed a relative decrease. Several factors can contribute to differences in specific findings across studies, including sampling methods and confounding factors. Our study used saliva samples, while others have more specific oral sampling such as plaque^[12]^. Antiretroviral therapy may affect the oral microbiome, though the specific effects remain unclear. Increased *Veillonella* was associated with INSTI-based antiretroviral therapy as well as an overweight BMI^[13]^. In our study population, the majority of participants were not on INSTI-based regimens (54.5%) and were not obese, suggesting that these factors do not explain our findings.

The positive correlations between *Veillonella parvula* IgG and microbial translocation markers (sCD14 and LBP) align with prior evidence suggesting systemic circulation of oral bacteria in inflammation-related disease states. In HIV, gut microbial translocation is a known cause of systemic inflammation^[5, 29-31]^. To date, no studies have examined oral microbial translocation in HIV. While studies have correlated oral bacteria with systemic immune biomarkers^[12, 15]^, none have investigated translocation mechanisms. In this study we demonstrated that *Veillonella parvula* may alter oral epithelial barrier permeability, promotes cytokine production and increases intermediate monocytes (**Figure 4**), indicating its capacity to enhance systemic inflammation in HIV.

There are some limitations to consider with this study. First, there was only limited oral health data available to include in the analysis. We did not have dental exams or records available. As other studies have shown the importance of oral health in oral microbiome composition in HIV^[32]^, this is an important limitation to note. Other important confounders including smoking, alcohol use, and substance use were all addressed in the analyses using a propensity-score based weighting method. Next, our *in vitro* studies were limited to representative bacterial species/strains that are widely available, which may differ from clinical species/strains *in vivo*. Further studies could include similar experiments using isolates cultures from clinical samples to confirm these findings. Our study identified potentially important associations between the oral microbiome and inflammation in HIV, and suggests a plausible mechanism through *Veillonella parvula* barrier disruption, translocation, and monocyte activation.

## Declaration of Interests

The authors have no conflicts of interest to declare.

## Acknowledgements

We would like to thank all mSTUDY participants for their generous participation in this work. This work was supported by funds from the California HIV/AIDS Research Program of the University of California, grant number BB19-LA-008 to J.A.F. This work was also supported in part by funds from NIH/NIDCR R01DE032624 to J.A.F, NIH/NIDA U01DA036267 to S.S. and P.M.G. Additional support was provided by the UCLA AIDS Institute and the UCLA-CDU Center for AIDS Research (NIH/NIAID P30AI152501). This publication includes data generated at the UC San Diego IGM Genomics Center utilizing an Illumina NovaSeq 6000 that was purchased with funding from a NIH SIG grant (S10 OD026929).

## Author Contributions

JAF, FL, NHT, GMA conceived of and designed the study. SS, PMG designed and provided study samples and data collection. KN, BY, JE, GDC performed experiments. JAF, FL performed data analysis. JAF wrote the original manuscript draft and all authors contributed to review and editing the manuscript.

## Notes

### Competing Interest Statement

The authors have declared no competing interest.

## REFERENCES

1. Deeks SG, Tracy R, Douek DC. Systemic effects of inflammation on health during chronic HIV infection. Immunity 2013; 39(4):633–645.

2. Neuhaus J, Jacobs DR, Jr., Baker JV, Calmy A, Duprez D, La Rosa A, et al. Markers of inflammation, coagulation, and renal function are elevated in adults with HIV infection. J Infect Dis 2010; 201(12):1788–1795.

3. Kuller LH, Tracy R, Belloso W, De Wit S, Drummond F, Lane HC, et al. Inflammatory and coagulation biomarkers and mortality in patients with HIV infection. PLoS Med 2008; 5(10):e203.

4. Sandler NG, Wand H, Roque A, Law M, Nason MC, Nixon DE, et al. Plasma levels of soluble CD14 independently predict mortality in HIV infection. J Infect Dis 2011; 203(6):780–790.

5. Brenchley JM, Price DA, Schacker TW, Asher TE, Silvestri G, Rao S, et al. Microbial translocation is a cause of systemic immune activation in chronic HIV infection. Nat Med 2006; 12(12):1365–1371.

6. Veazey RS, DeMaria M, Chalifoux LV, Shvetz DE, Pauley DR, Knight HL, et al. Gastrointestinal tract as a major site of CD4+ T cell depletion and viral replication in SIV infection. Science 1998; 280(5362):427–431.

7. Olsson J, Poles M, Spetz AL, Elliott J, Hultin L, Giorgi J, et al. Human immunodeficiency virus type 1 infection is associated with significant mucosal inflammation characterized by increased expression of CCR5, CXCR4, and beta-chemokines. J Infect Dis 2000; 182(6):1625–1635.

8. Vujkovic-Cvijin I, Dunham RM, Iwai S, Maher MC, Albright RG, Broadhurst MJ, et al. Dysbiosis of the gut microbiota is associated with HIV disease progression and tryptophan catabolism. Sci Transl Med 2013; 5(193):193ra191.

9. Dang AT, Cotton S, Sankaran-Walters S, Li CS, Lee CY, Dandekar S, et al. Evidence of an increased pathogenic footprint in the lingual microbiome of untreated HIV infected patients. BMC Microbiol 2012; 12:153.

10. Saxena D, Li Y, Devota A, Pushalkar S, Abrams W, Barber C, et al. Modulation of the orodigestive tract microbiome in HIV-infected patients. Oral Dis 2016; 22 Suppl 1:73–78.

11. Beck JM, Schloss PD, Venkataraman A, Twigg H, 3rd, Jablonski KA, Bushman FD, et al. Multicenter Comparison of Lung and Oral Microbiomes of HIV-infected and HIV-uninfected Individuals. Am J Respir Crit Care Med 2015; 192(11):1335–1344.

12. Annavajhala MK, Khan SD, Sullivan SB, Shah J, Pass L, Kister K, et al. Oral and Gut Microbial Diversity and Immune Regulation in Patients with HIV on Antiretroviral Therapy. mSphere 2020; 5(1):e00798–00719.

13. Narayanan A, Kieri O, Vesterbacka J, Manoharan L, Chen P, Ghorbani M, et al. Exploring the interplay between antiretroviral therapy and the gut-oral microbiome axis in people living with HIV. Sci Rep 2024; 14(1):17820.

14. Li J, Chang S, Guo H, Ji Y, Jiang H, Ruan L, et al. Altered Salivary Microbiome in the Early Stage of HIV Infections among Young Chinese Men Who Have Sex with Men (MSM). Pathogens 2020; 9(11).

15. Yang L, Dunlap DG, Qin S, Fitch A, Li K, Koch CD, et al. Alterations in Oral Microbiota in HIV Are Related to Decreased Pulmonary Function. Am J Respir Crit Care Med 2020; 201(4):445–457.

16. Noguera-Julian M, Guillen Y, Peterson J, Reznik D, Harris EV, Joseph SJ, et al. Oral microbiome in HIV-associated periodontitis. Medicine (Baltimore) 2017; 96(12):e5821.

17. Li Y, Saxena D, Chen Z, Liu G, Abrams WR, Phelan JA, et al. HIV infection and microbial diversity in saliva. J Clin Microbiol 2014; 52(5):1400–1411.

18. Li S, Zhu J, Su B, Wei H, Chen F, Liu H, et al. Alteration in Oral Microbiome Among Men Who Have Sex With Men With Acute and Chronic HIV Infection on Antiretroviral Therapy. Front Cell Infect Microbiol 2021; 11:695515.

19. Presti RM, Handley SA, Droit L, Ghannoum M, Jacobson M, Shiboski CH, et al. Alterations in the oral microbiome in HIV-infected participants after antiretroviral therapy administration are influenced by immune status. Aids 2018; 32(10):1279–1287.

20. Peng X, Cheng L, You Y, Tang C, Ren B, Li Y, et al. Oral microbiota in human systematic diseases. Int J Oral Sci 2022; 14(1):14.

21. Chhibber-Goel J, Singhal V, Bhowmik D, Vivek R, Parakh N, Bhargava B, et al. Linkages between oral commensal bacteria and atherosclerotic plaques in coronary artery disease patients. NPJ Biofilms Microbiomes 2016; 2:7.

22. Rojas-Tapias DF, Brown EM, Temple ER, Onyekaba MA, Mohamed AMT, Duncan K, et al. Inflammation-associated nitrate facilitates ectopic colonization of oral bacterium Veillonella parvula in the intestine. Nat Microbiol 2022; 7(10):1673–1685.

23. Cook RR, Fulcher JA, Tobin NH, Li F, Lee D, Javanbakht M, et al. Effects of HIV viremia on the gastrointestinal microbiome of young MSM. Aids 2019; 33(5):793–804.

24. Cook RR, Fulcher JA, Tobin NH, Li F, Lee D, Woodward C, et al. Combined effects of HIV and obesity on the gastrointestinal microbiome of young men who have sex with men. HIV Med 2020; 21(6):365–377.

25. Marotz C, Cavagnero Kellen J, Song Se J, McDonald D, Wandro S, Humphrey G, et al. Evaluation of the Effect of Storage Methods on Fecal, Saliva, and Skin Microbiome Composition. mSystems 2021; 6(2):10.1128/msystems.01329-01320.

26. Yang X, Fox A, Powell RLR. Qualitative immunoassay for the detection of anti-SARS-COV-2 spike antibody in human milk samples. STAR Protocols 2022; 3(1):101203.

27. Fulcher JA, Hussain SK, Cook R, Li F, Tobin NH, Ragsdale A, et al. Effects of Substance Use and Sex Practices on the Intestinal Microbiome During HIV-1 Infection. J Infect Dis 2018; 218(10):1560–1570.

28. Jacobsen J, van Deurs B, Pedersen M, Rassing MR. TR146 cells grown on filters as a model for human buccal epithelium: I. Morphology, growth, barrier properties, and permeability. International Journal of Pharmaceutics 1995; 125(2):165–184.

29. Dillon SM, Lee EJ, Kotter CV, Austin GL, Dong Z, Hecht DK, et al. An altered intestinal mucosal microbiome in HIV-1 infection is associated with mucosal and systemic immune activation and endotoxemia. Mucosal Immunol 2014; 7(4):983–994.

30. Jiang W, Lederman MM, Hunt P, Sieg SF, Haley K, Rodriguez B, et al. Plasma levels of bacterial DNA correlate with immune activation and the magnitude of immune restoration in persons with antiretroviral-treated HIV infection. J Infect Dis 2009; 199(8):1177–1185.

31. Klase Z, Ortiz A, Deleage C, Mudd JC, Quiñones M, Schwartzman E, et al. Dysbiotic bacteria translocate in progressive SIV infection. Mucosal Immunol 2015; 8(5):1009–1020.

32. Griffen AL, Thompson ZA, Beall CJ, Lilly EA, Granada C, Treas KD, et al. Significant effect of HIV/HAART on oral microbiota using multivariate analysis. Sci Rep 2019; 9(1):19946.

