## Supplemental Figure 1 for "Comparison of oral and gut microbiome highlights role of oral bacteria in systemic inflammation in HIV"

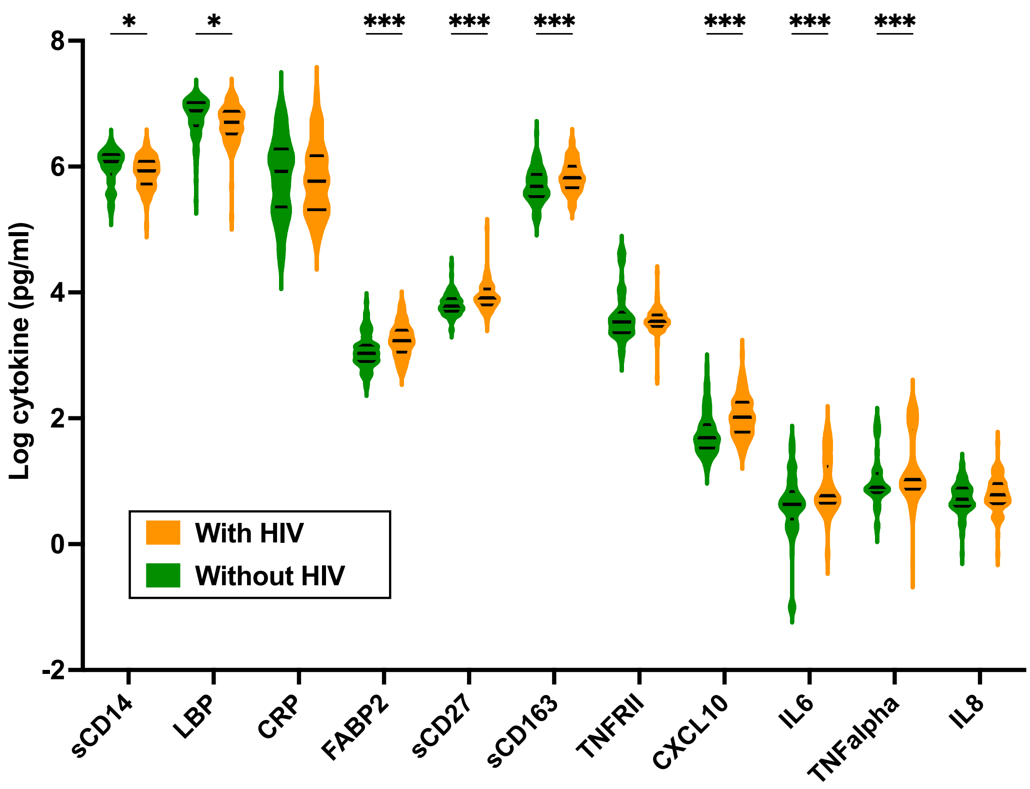


**Supplementary Figure 1**. Plasma cytokines/biomarkers by HIV groups. Middle lines denote median with top/bottom lines quartiles. *p<0.05; *** p<0.001 by unpaired t-tests.
